# PRDM3 attenuates pancreatitis and pancreatic tumorigenesis by regulating inflammatory response

**DOI:** 10.1101/2019.12.18.880823

**Authors:** Jie Ye, Anpei Huang, Haitao Wang, Anni M.Y. Zhang, Xiaojun Huang, Qingping Lan, Tomohiko Sato, Susumu Goyama, Mineo Kurokawa, Chuxia Deng, Maike Sander, David F. Schaeffer, Wen Li, Janel L. Kopp, Ruiyu Xie

## Abstract

Pancreatic ductal adenocarcinoma (PDAC) is associated with metaplastic changes in the pancreas but the transcriptional program underlying these changes is incompletely understood. The zinc finger transcription factor, PRDM3, is lowly expressed in normal pancreatic acini and its expression increases during tumorigenesis. Although PRDM3 promotes proliferation and migration of PDAC cell lines, the role of PRDM3 during tumor initiation from pancreatic acinar cells *in vivo* is unclear. In this study, we showed that high levels of PRDM3 expression in human pancreas was associated with pancreatitis, and well-differentiated but not poorly differentiated carcinoma. We examined PRDM3 function in pancreatic acinar cells during tumor formation and pancreatitis by inactivating *Prdm3* using a conditional allele (*Ptf1a^CreER^;Prdm3^flox/flox^*mice) in the context of oncogenic Kras expression and supraphysiological cerulein injections, respectively. In *Prdm3*-deficient mice, Kras^G12D^-driven preneoplastic lesions were more abundant and progressed to high-grade precancerous lesions more rapidly. This is consistent with our observations that low levels of PRDM3 in human PDAC was correlated significantly with poorer survival in patient. Moreover, loss of *Prdm3* in acinar cells elevated exocrine injury, enhanced immune cell activation and infiltration, and greatly increased acinar-to-ductal cell reprogramming upon cerulein-induced pancreatitis. Whole transcriptome analyses of *Prdm3* knockout acini revealed that pathways involved in inflammatory response and Hif-1 signaling were significantly upregulated in *Prdm3*-depleted acinar cells. Taken together, our results suggest that Prdm3 favors the maintenance of acinar cell homeostasis through modulation of their response to inflammation and oncogenic Kras activation, and thus plays a previously unexpected suppressive role during PDAC initiation.

## Introduction

Pancreatic ductal adenocarcinoma (PDAC) is the most common malignancy in pancreas and the third leading cause of cancer-related deaths in US^1^. Emerging evidence suggests that pancreatic acinar cells can acquire ductal cell-like characteristics and down-regulate genes maintaining acinar cell identify, also known as acinar-to-ductal metaplasia (ADM). This process is reversible because once the injury is resolved the acinar-cell-derived ductal-like cells can revert back to acinar cells^2, 3^. However, in the presence of additional stresses, such as a *Kras^G12D^*mutation, ADM cannot be reversed and cells are “locked” into a transdifferentiated state before converting to precancerous pancreatic intraepithelial neoplasia (PanIN) lesions and subsequently invasive PDAC^4–6^. In mice, pancreatic tumorigenesis is dramatically hastened by the presence of pancreatitis^7, 8^, while, in humans, induction of chronic inflammation is a common character in many known risk factors for pancreatic cancer including diabetes, pancreatitis, alcohol consumption and tobacco use^9^. However, the complete transcriptional program that regulates the interconversion of acinar cells to ductal-like cells and vice versa, and the role of these events in the context of tumorigenesis are still unclear.

PRDM3 is a nuclear transcription factor involved in many biological processes including hematopoiesis, development, cell differentiation and apoptosis^10^. PRDM3 belongs to the positive regulatory domain (PRDM) family proteins, which are characterized by an N-terminal PR (PRDI-BF1-RIZ1 homologous) domain followed by an array of C2H2 zinc finger motifs for sequence-specific DNA binding and a C-terminal binding protein (CtBP)-binding domain for protein-protein interactions^11^. PRDM3 is necessary for the maintenance of hematopoietic stem cells^12, 13^. A recent study has reported that PRDM3 is weakly expressed in normal pancreatic acinar cells and upregulated in many PDAC precursor lesions and PDAC^14^. Using siRNA-mediated knockdown of PRDM3 in PK-8 pancreatic cancer cells, Tanaka and colleagues also showed that PRDM3 promotes pancreatic cancer cell proliferation and migration through the inhibition of a KRAS suppressor miR-96^14^. Despite this characterization of the effects of PRDM3 inhibition in pancreatic tumor cells *ex vivo*, the role of PRDM3 during tumor initiation from acinar cells *in vivo* is unclear.

Here, we used a CreER-inducible mouse model to genetically delete *Prdm3* specifically in adult acinar cells to examine the functional role of PRDM3 in pancreatic carcinogenesis. Our results show that *Prdm3* deficiency potentiates inflammation, promotes tumour initiation and dramatically accelerates malignant progression, which is consistent with our findings indicating that PRDM3 loss is significantly associated with poorer survival in patients with PDAC. We further demonstrate that PRDM3 is important to suppress the expression of genes involved in inflammatory response in pancreatic acinar cells. These findings suggest an inhibitory role of PRDM3 in pancreatic tumorigenesis. Future development of drugs that target PRDM3 might yield novel approaches to benefit the treatment of PDAC.

## Results

### PRDM3 is upregulated in pancreatitis as well as well-differentiated PDAC and its high expression is associated with better survival in patients with PDAC

We first characterized the expression of PRDM3 and its relevance to pancreatic cancer prognosis by analyzing a cohort of 94 patients who were diagnosed with PDAC and received surgical resection without preoperative chemotherapy. We found that PRDM3 was strongly expressed in precancerous PanIN lesions (**Fig. 1c-c’)** and well-differentiated PDAC from patients (**Fig. 1d-d’)**, while moderately to poorly differentiated cancer cells showed little to no staining of PRDM3 **(Fig. 1e-e**’ **and f-f’, Table 1)**. Our observation of heterogeneous expression of PRDM3 in PDAC was supported by recent study demonstrating that PRDM3 is selectively expressed in low-grade PDAC cells featured differentiated epithelia, but not high-grade cells showed fibroblastoid morphology^15^. We performed subsequent overall survival and disease-free survival analyses with these 94 PDAC patients and found that patients with high levels of PRDM3 lived significantly longer than those with low levels of PRDM3 (Overall survival: 16.03 months vs 9.33 months; Disease-free survival: 12.37 months vs 7.4 months) **(Fig. 1g)**. Our clinical relevance analysis clearly revealed that a better survival in patient with PDAC was associated with high levels of PRDM3 expression, but not with age, gender, tumor size, location, TNM (tumour-node-metastasis), or CA19-9 (**Table 1**). Given that pancreatitis is a well-described risk factor for PDAC development, we also analyzed pancreatic tissue from 22 patients with chronic pancreatitis. We found a dramatically increase of PRDM3 protein levels in inflamed tissues compared with normal pancreas **(Fig. 1a-a’ and b-b’; Supplementary Table 1)**. Similarly, administration of supraphysiologic concentrations of a cholecystokinin ortholog, cerulein, in mice resulted in acute pancreatitis and Prdm3 upregulation in murine acinar cells **(Supplementary Fig. 1a)**. Consistent with findings from previous reports^14^, we also observed strong expression of Prdm3 in the precursor lesions of PDAC including ADM, low-grade PanINs, and high-grade PanINs found in pancreata from mice expressing oncogenic *Kras* in pancreatic acinar cells (*Ptf1a^CreER^;Kras^G12D^*) **(Supplementary Fig. 1b)**. Together, our results demonstrated that elevated levels of PRDM3 are associated with inflamed pancreatic epithelia and well-differentiated pancreatic lesions, while low levels of PRDM3 are associated with poorly differentiated carcinoma and a worse prognostic outcome in patients with PDAC.

**Fig. 1.**
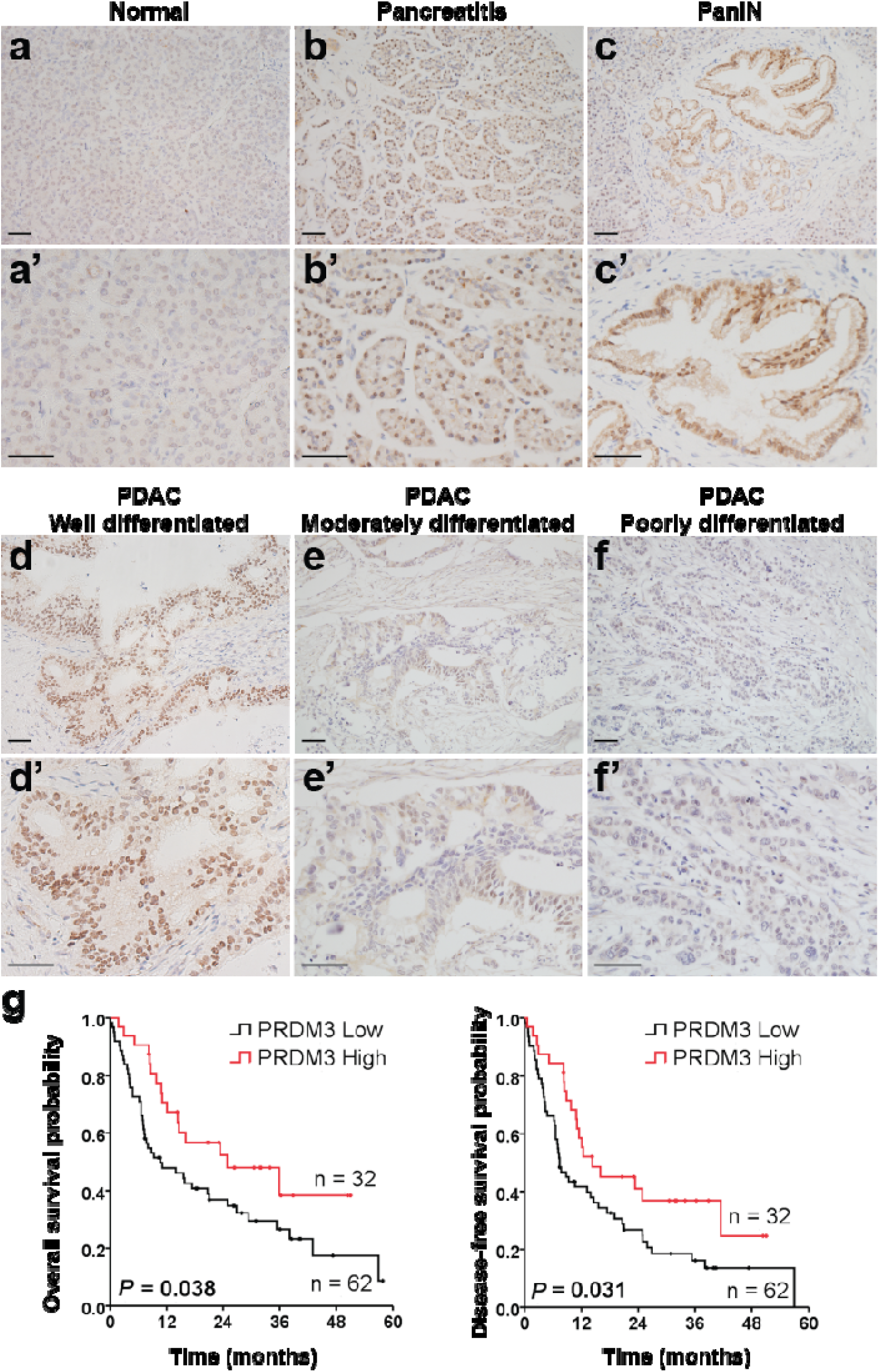
PRDM3 underexpression is associated with poorly differentiated tumors and a worse prognostic outcome in patients with PDAC. Prdm3 immunostaining in normal pancreatic tissue **(a and a’)**, pancreatitis **(b and b’)**, PanIN **(c and c’)**, well-differentiated **(d and d’)**, moderately differentiated **(e and e’)** and poorly differentiated **(f and f’)** PDAC. **(g)** The overall survival probability and disease-free survival probability were compared between low (n=32, staining index ≤ 6) and high (n=62, staining index > 6) levels of PRDM3 expression in a cohort of 94 PDAC patients after surgical resection. *p*-values were calculated based on log-rank test. Scale: 50 μm.

**Table 1.**
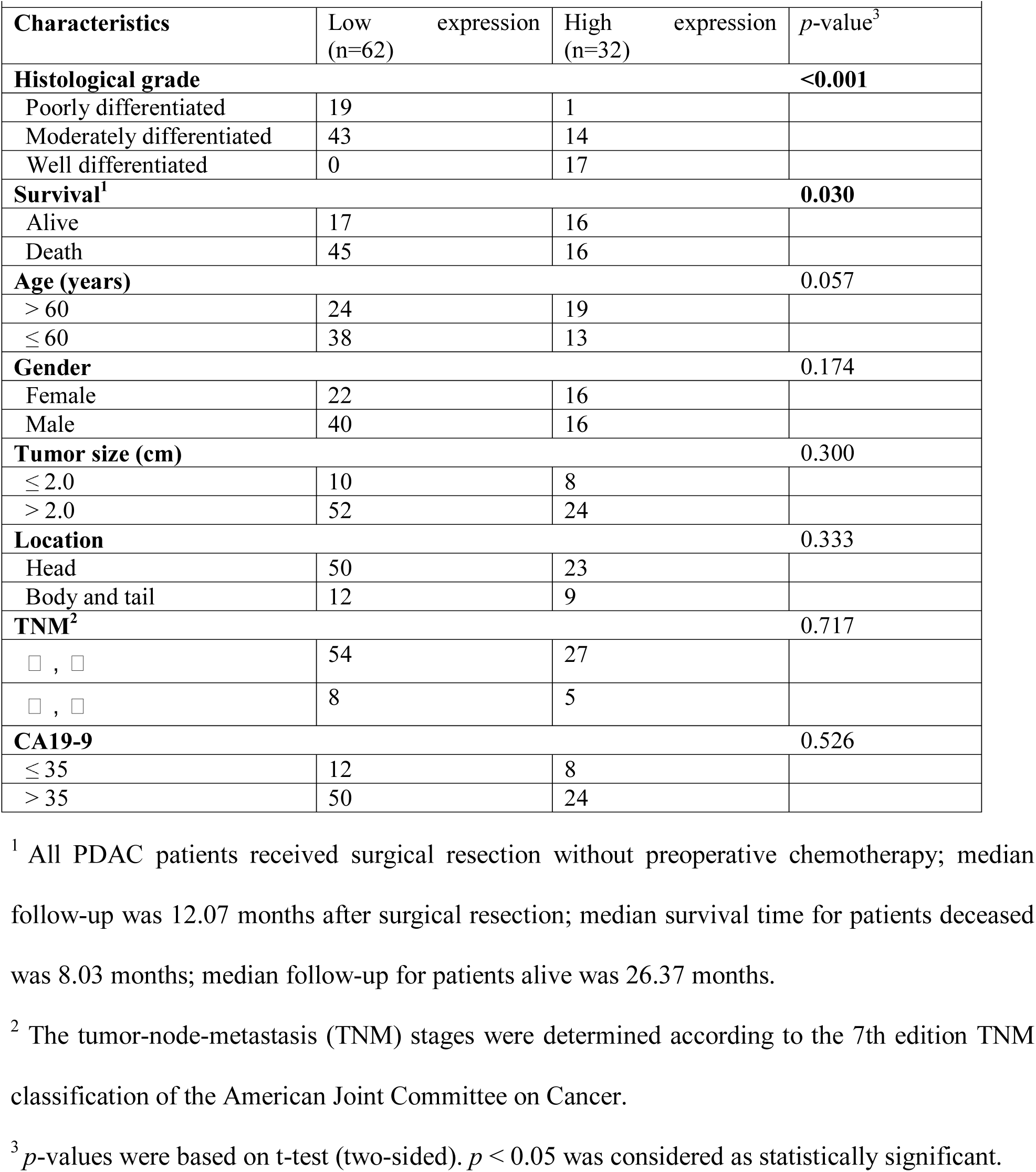
Association between expression levels of PRDM3 and clinical relevance in patients with PDAC (n=94)

### Ablation of *Prdm3* enhances Kras^G12D^-stimulated PDAC initiation and progression

To determine whether Prdm3 is functionally important for pancreatic carcinogenesis *in vivo*, we applied a genetic strategy to induce expression of oncogenic *Kras^G12D^* and deletion of *Prdm3* in adult acinar cells, simultaneously. Cre-mediated recombination was induced in pancreatic acinar cells using the tamoxifen-inducible *Ptf1a^CreER^* allele^16^. The Mds1 and Evi1 complex locus (*Mecom*) encodes a full-length isoform of *Prdm3*, and a shorter isoform lacking the N-terminus PR domain. We therefore used a *Prdm3^flox^* mouse which harbors two *LoxP* sites flanking exon 4, the first shared exon in the long- and short-isoform of *Prdm3*^12^, to completely eliminate *Prdm3* in pancreatic acinar cells upon tamoxifen induced recombination. By combining the *Kras^LSL-G12D^*allele^17^ with the *Ptf1a^CreER^*allele with/without the *Prdm3^flox^* allele, we generated control *Ptf1a^CreER^;Kras^G12D^*(*Kras^G12D^*) mice, as well as *Ptf1a^CreER^;Kras^G12D^;Prdm3^flox/flox^*(*Kras^G12D^-Prdm3^ΔAcinar^*) mice **(Supplementary Fig. 2a)**.

To initiate recombination, we injected mice with tamoxifen at 4 to 5 weeks of age and analyzed pancreata at 4- and 6-weeks post-injection **(Fig. 2a)**. Comparison of Prdm3 expression in *Kras^G12D^-Prdm3^ΔAcinar^* and *Kras^G12D^* mice after tamoxifen administration showed an almost complete loss of Prdm3 protein in *Kras^G12D^-Prdm3^ΔAcinar^* acinar cells **(Supplementary Fig. 2b)**. Four weeks after tamoxifen-mediated recombination, small areas of ADM and occasional low-grade PanINs were observed in the control *Kras^G12D^* mice. In contrast, *Kras^G12D^-Prdm3^ΔAcinar^* mice, with loss of Prdm3 proteins in pancreatic acinar cells, exhibited more cuboidal to columnar duct-like structures with enlarged lumens **(Supplementary Fig. 2c)**. Quantification of the number of PanINs revealed that preneoplastic lesions arising in *Kras^G12D^-Prdm3^ΔAcinar^* mice increased significantly compared to *Kras^G12D^* mice **(Fig. 2b)**. More intriguingly, we observed 4 out of 5 *Kras^G12D^-Prdm3^ΔAcinar^* mice developed high-grade PanINs at 4 weeks post-tamoxifen injection **(Fig. 2c)**, while no high-grade lesions were found in *Kras^G12D^* mice even at 3 months of age (data not shown). Consistent with these findings, at 6 weeks post-tamoxifen injection *Kras^G12D^-Prdm3^ΔAcinar^* mice exhibited higher number of lesions with histological and molecular characteristics of PanINs indicated by the expression of Cytokeratin 19 (CK19) and Mucin 5AC (Muc5AC) **(Fig. 2d)**. Together, these data suggest that deletion of *Prdm3* promotes ADM and PanIN formation in the presence of oncogenic *Kras* expression.

**Fig. 2.**
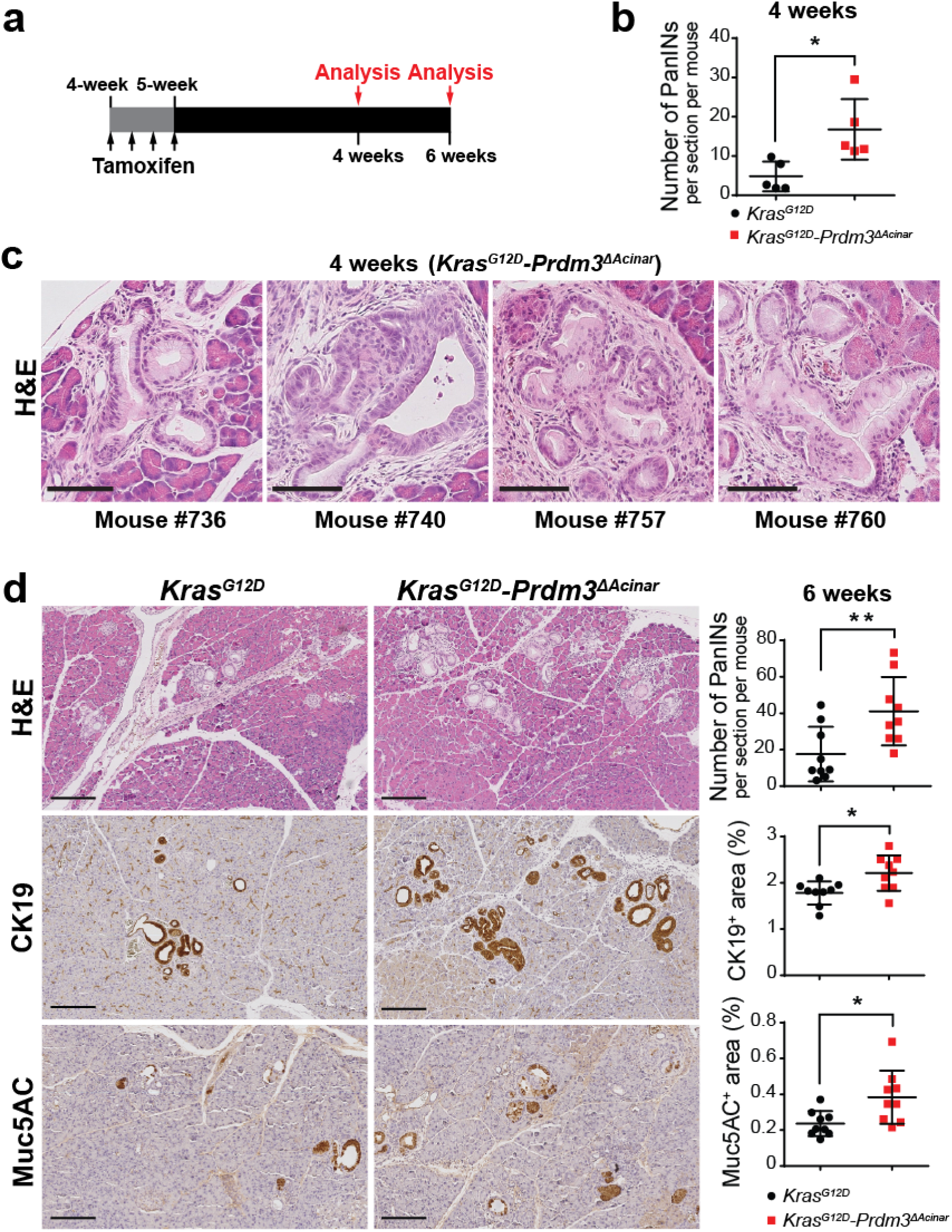
Loss of prdm3 promotes acinar-to-ductal metaplasia and PanIN lesions formation. **(a)** *Ptf1a^CreER^;Kras^G12D^*and *Ptf1a^CreER^;Kras^G12D^;Prdm3^flox/flox^*mice at 4-5 weeks of age were injected 4 times on alternating days with tamoxifen. Recombined mice were analyzed at 4 and 6 weeks post tamoxifen injection. **(b)** The number of PanINs per section for *Kras^G12D^* (n=5) vs. *Kras^G12D^-Prdm3^ΔAcinar^* (n=5) mice at 4 weeks post tamoxifen injection. **(c)** Representative images of high-grade PanINs in *Kras^G12D^-Prdm3^ΔAcinar^* mice at 4 weeks after tamoxifen injection. **(d)** Hematoxylin-eosin staining (H&E), immunohistochemistry staining for ductal marker Cytokeratin 19 (CK19) and mucinous maker Muc5AC. Quantification of the number of PanINs, as well as the respective percent of pancreatic area that is CK19^+^ and Muc5AC^+^, at 6 weeks post tamoxifen injection in *Kras^G12D^* (n=9) vs. *Kras^G12D^-Prdm3^ΔAcinar^* (n=9) mice. Data show mean ± SD. \**p* < 0.05, \*\**p* < 0.01. Scale: 200 μm.

We next examined whether loss of *Prdm3* promotes neoplastic progression. To accelerate the formation of invasive lesions, we induced cerulein-mediated acute pancreatitis in cooperation with acinar-cell-specific activation of oncogenic Kras as previously described^4^. One week after tamoxifen administration, *Kras^G12D^-Prdm3^ΔAcinar^* and *Kras^G12D^* mice were injected hourly with 50 μg/kg cerulein over 6 hours on alternating days. The pancreata were harvested at 21 days post-cerulein injection **(Fig. 3a)**. Pancreata from *Kras^G12D^-Prdm3^ΔAcinar^* mice had full spectrum of precursor lesions including low-grade PanINs, high-grade PanINs and ductal carcinoma *in situ* **(Supplementary Fig. 3a)**, which had lost Prdm3 staining **(Supplementary Fig. 3b)**. The control *Kras^G12D^* mice had less evidence of tumorigenesis compared with *Kras^G12D^-Prdm3^ΔAcinar^* mice. Specifically, in *Kras^G12D^-Prdm3^ΔAcinar^* mice, the ratio of pancreas-to-body weight increased **(Fig. 3b)**; the number of Cpa1^+^acinar cells decreased dramatically and CK19^+^ duct-like cells increased **(Fig. 3c)**; a higher percentage of the pancreas was replaced by acidic mucin content indicated by Alcian Blue staining **(Fig. 3c)**; and the number of high-grade PanIN significantly elevated **(Fig. 3e)**. Moreover, we consistently observed tumor budding associated with many high-grade neoplastic lesions in *Kras^G12D^-Prdm3^ΔAcinar^* mice at 21 days post-cerulein injection **(Fig. 3d)**. Tumor budding is a strong prognostic indicator of aggressive tumor behavior, which is defined as the presence of single cells or clusters of less than five tumor cells scattered in the stroma^18^. In contrast to *Kras^G12D^-Prdm3^ΔAcinar^* mice, tumor budding was rarely observed in *Kras^G12D^* mice, suggesting that loss of *Prdm3* accelerates pancreatic cancer formation in *Kras^G12D^*-expressing mice. Together, our data suggest that progression of low-grade precursor lesions to high-grade PanIN is more rapid in the absence of Prdm3.

**Fig. 3.**
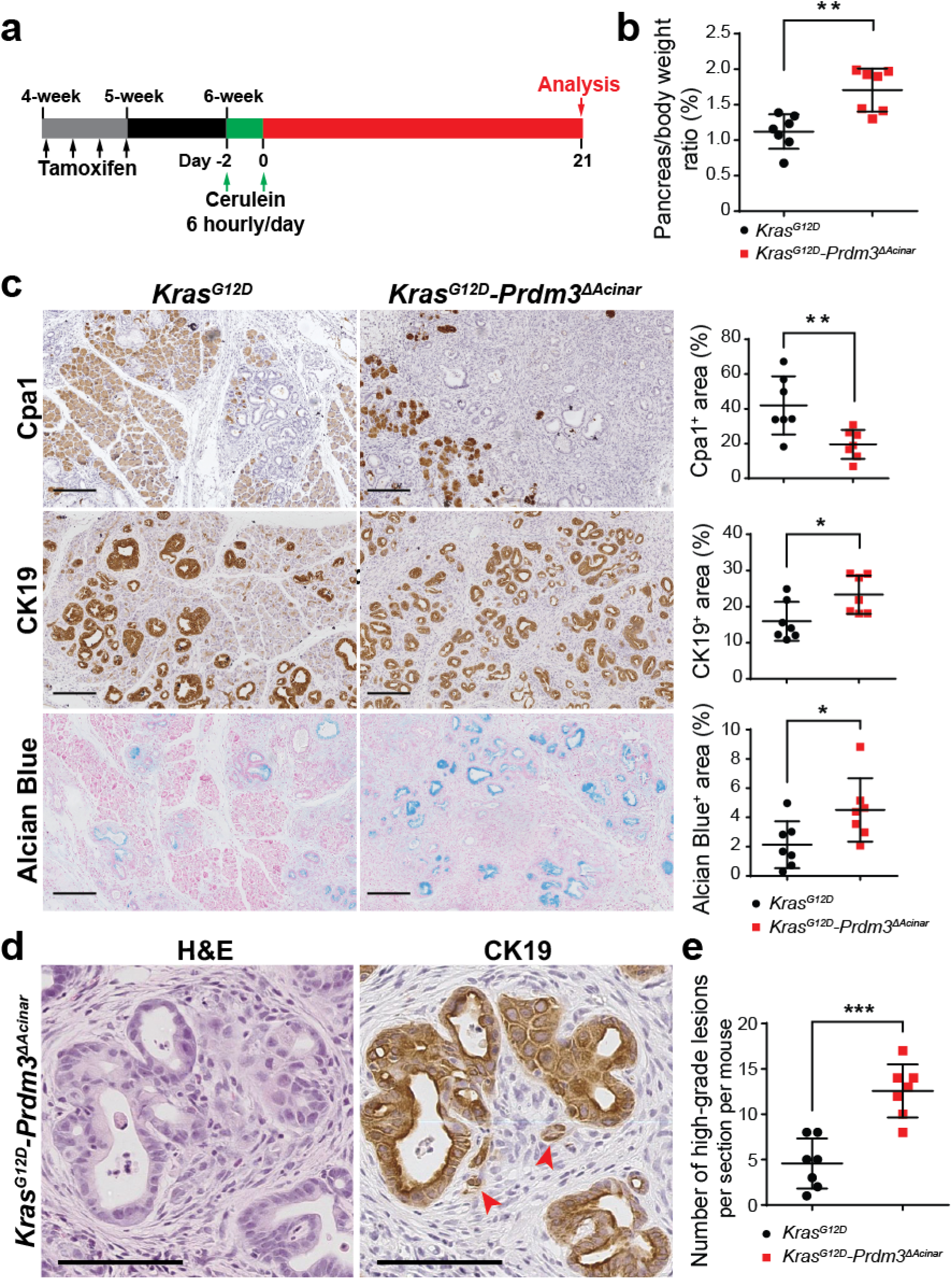
Inhibition of prdm3 accelerates Kras^G12D^-driven neoplastic transformation in response to pancreatitis. **(a)** Schematic illustration showing experimental design of cerulein-induced acute pancreatitis in cooperation with activation of oncogenic *Kras* in *Ptf1a*-expressing cells. *Ptf1a^CreER^;Kras^G12D^*(n=7) and *Ptf1a^CreER^;Kras^G12D^;Prdm3^flox/flox^*(n=7) mice at 4-5 weeks of age were injected 4 times on alternating days with tamoxifen. One week after the last tamoxifen injection, mice were subjected to cerulein (50 μg/kg) injection at hourly intervals over 6 hours on alternating days separated by 24 hours and analyzed at 21 days post cerulein injection. **(b)** Quantification of relative pancreas mass measured as per cent of pancreas weight over body weight in *Kras^G12D^*vs *Kras^G12D^-Prdm3^ΔAcinar^* mice. **(c)** Hematoxylin-eosin staining (H&E) and immunohistochemistry for Cpa1, Cytokeratin 19 (CK19) and Alcian Blue of pancreata from *Kras^G12D^*and *Kras^G12D^-Prdm3^ΔAcinar^* mice. Quantification of the percent of pancreatic area that is Cpa1^+^, CK19^+^ and Alcian Blue^+^ in *Kras^G12D^* vs. *Kras^G12D^-Prdm3^ΔAcinar^* mice. **(d)** Representative images of tumor budding in *Kras^G12D^-Prdm3^ΔAcinar^* mice indicated by red arrowheads. Immunohistochemistry staining CK19 strongly suggests invasive high-grade neoplasia. **(e)** Number of high-grade PanINs per section for each genotype. Data show mean ± SD. \**p* < 0.05, \*\**p* < 0.01, *** *p* < 0.001. Scale: 200 μm (c) and 100 μm (d).

### Loss of *Prdm3* in acinar cells enhances pancreatitis

Given that inflammation promotes cancer formation and Prdm3 was significantly increased in humans with pancreatitis, we further determined whether Prdm3 modulated the inflammatory response of the pancreas in *Ptf1a^CreER^;Prdm3^flox/flox^*mice. Mice injected with tamoxifen, but lacking the *Prdm3^flox^*allele, were used as controls (*Ptf1a^CreER^*mice). Seven days after tamoxifen administration, the Prdm3 protein was absent in more than 80% of acinar cells in *Prdm3^ΔAcinar^* mice, indicating efficient and specific deletion of Prdm3 **(Supplementary Fig. 4a)**. Acute pancreatitis was induced with intraperitoneal injection of cerulein, as described previously^19^ **(Fig. 4a)**. Specifically, 7 days after the last tamoxifen injection, *Prdm3^ΔAcinar^* and control (*Ptf1a^CreER^*) mice were injected with 50 μg/kg cerulein at hourly intervals for 8 hours. Histological assessment of pancreata from mice 3 hours after the last cerulein injection demonstrated exaggerated interstitial edema, cytoplasmic vacuolization and immune cell infiltration in *Prdm3^ΔAcinar^* mice compared with control mice **(Fig. 4b)**. Consistently, examination of neutrophil infiltration demonstrated a substantial increase in the number of Ly6B.2^+^ cells in cerulein-treated *Prdm3^ΔAcinar^* pancreata **(Fig. 4b)**. Elevated blood amylase levels are an indicator of pancreatitis. Consistent with the neutrophil infiltration in the pancreas, we observed significantly higher levels of serum amylase in *Prdm3^ΔAcinar^* mice compared to controls **(Fig. 4c)**. To determine whether the expression of inflammatory cytokines is increased, we harvested RNA from pancreata of *Prdm3^ΔAcinar^* and control mice to performe quantitative RT-PCR. As expected, expression of inflammatory cytokines, including *Il-6*, *Cxcl-1*, *Cxcl-10*, *Ccl2*, and *Ccl20*, increased significantly in *Prdm3*-deleted compared to control pancreata **(Supplementary Fig. 4b)**. We further examined pancreata harvested from *Prdm3^ΔAcinar^* and control mice 48 hours after two series of 8-hourly injection of cerulein **(Fig. 4a)**. Characterization by immunohistochemistry for the ductal marker CK19 confirmed that acini undergoing acinar-to-ductal metaplasia increased dramatically in *Prdm3^ΔAcinar^* mice **(Fig. 4d)**. These results illustrate that acinar-cell-specific ablation of *Prdm3* augments the severity of pancreatitis, suggesting a specific role of Prdm3 as a modulator of inflammatory response in the pancreas.

**Fig. 4.**
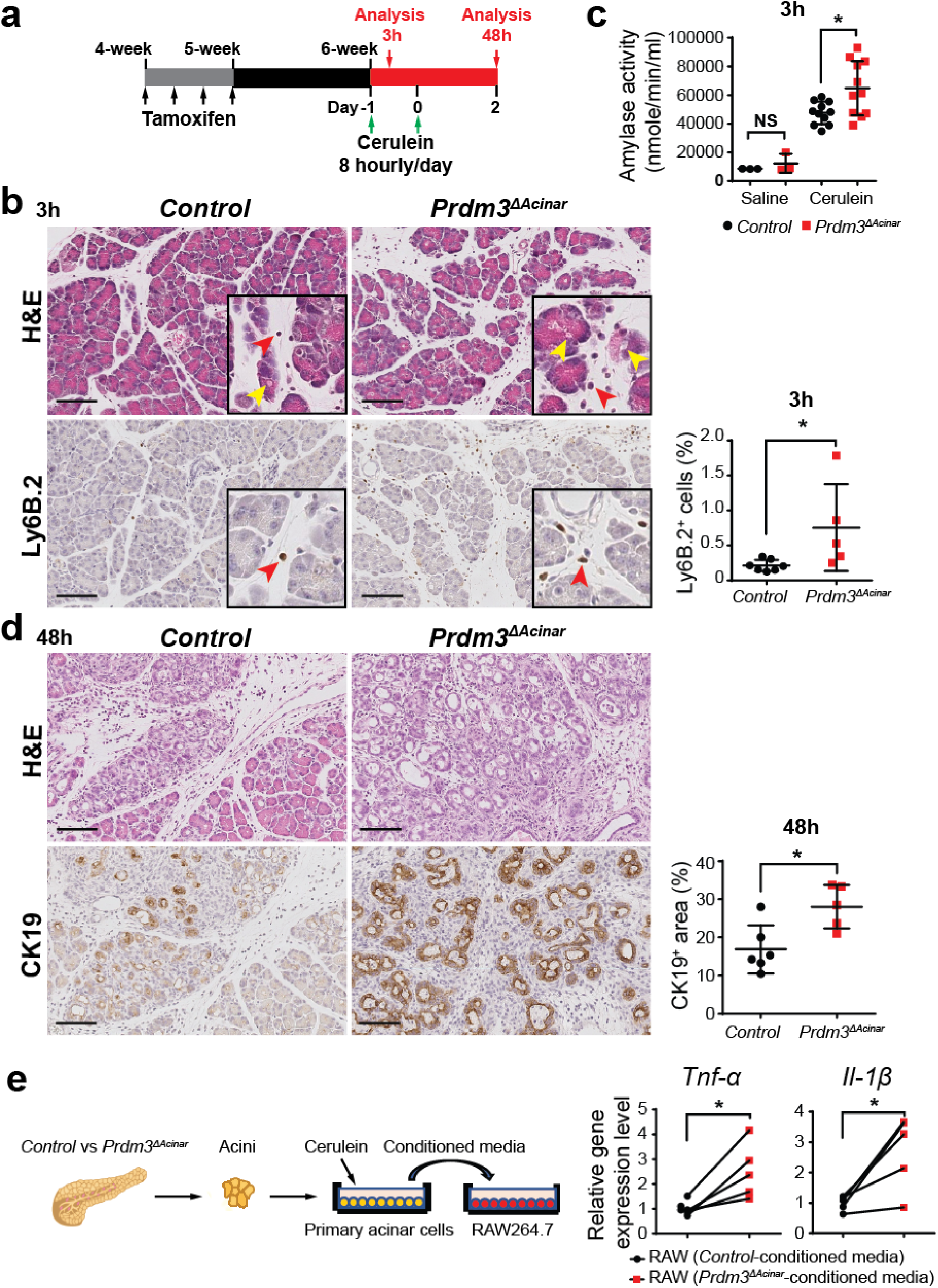
Prdm3 deletion in pancreatic acinar cells exaggerates cerulein-induced pancreatitis. **(a)** Schematic illustration of experimental design. *Ptf1a^CreER^;Prdm3^flox/flox^*mice at 4-5 weeks of age were injected 4 times on alternating days with tamoxifen. One week after the last tamoxifen injection, mice were subjected to cerulein (50 μg/kg) injection at hourly intervals for 8 hours per day for two consecutive days and analyzed at the indicated time points. **(b)** Histologic characterization was determined with hematoxylin-eosin staining (H&E). Immunohistochemistry neutrophil marker Ly6B.2 with quantitation of the percent of all cells that are Ly6B.2^+^ in *Ptf1a^CreER^* (control, n=7) and *Prdm3^ΔAcinar^* (n=5) pancreata. Vacuole is indicated by yellow arrowheads, and infiltrated immune cells are indicated by red arrowheads. **(c)** Serum amylase levels of cerulein-treated control (n=11) vs *Prdm3^ΔAcinar^* (n=11) mice and saline-treated control (n=3) vs *Prdm3^ΔAcinar^* (n=3). **(d)** Representative images of control and *Prdm3^ΔAcinar^* pancreata stained with H&E and ductal marker Cytokeratin 19 (CK19). Quantification of the respective percent of all cells that are CK19^+^ from control (n=6) vs *Prdm3^ΔAcinar^* (n=5) pancreata. **(e)** Primary acinar cells were isolated from control (n=5) vs *Prdm3^ΔAcinar^* (n=5) mice and cultured in Waymouth media in the presence of 10 nM cerulein for 6 hours. Macrophage cells Raw264.7 were treated with the above acinar-cell-conditioned media for 16 hours and collected for quantitative real-time PCR for the indicated cytokines. Each set of connected dots in (e) depicts a biological replicate (n=5 experiments). Data show mean ± SD. \**p* < 0.05. Scale: 100 μm.

Injured acinar cells initiate inflammatory responses by releasing proinflammatory cytokines, digestive enzymes, and nuclear damage-associated molecular patterns molecules, which attract and activate immune cell to exacerbate tissue injury^20, 21^. It has been demonstrated that pancreatic acinar cells undergo necrosis when exposed to supraphysiological cerulein *in vitro*^22^. To investigate whether loss of *Prdm3* in acinar cells can enhance the activation of inflammatory cells upon cerulein treatment, we exposed the murine macrophage cells, Raw246.7, to the conditioned media from *Prdm3^ΔAcinar^* or *Ptf1a^CreER^*control acini treated with cerulein (**Fig. 4e**). Briefly, acinar cells were isolated from a cohort of *Prdm3^ΔAcinar^* and control mice. Isolated primary acinar cells were allowed to recover in oxygenated medium for 2 hours and subsequently cultured in the presence of 10 nM cerulein for 6 hours. Raw246.7 macrophage cells were incubated in this acinar cell conditioned media for 16 hours. Activation of macrophages was determined by the expression of inflammatory cytokines. We found that the relative expression of inflammatory cytokines *Tnf-*α and *Il-1*β was significantly higher in Raw246.7 incubated with *Prdm3^ΔAcinar^* conditioned media (**Fig. 4e**), suggesting that more proinflammatory factors were released from *Prdm3*-deficient cells than controls to stimulate macrophage activation. These results support our findings that *Prdm3^ΔAcinar^* mice are more susceptible to cerulein-induced injury in part by contributing to increased immune cells activation.

### Loss of Prdm3 activates inflammatory response and Hif-1 signaling pathways

To further examine the effect of *Prdm3* deletion on acinar cell homeostasis, we performed transcriptome analyses of primary acinar cells isolated from *Prdm3^ΔAcinar^* and control mice. *Ptf1a^CreER^;Prdm3^flox/flox^*and control *Ptf1a^CreER^* mice were injected with four doses of tamoxifen at 4 to 5 weeks of age to induce recombination prior to acini isolation. One week after the last tamoxifen injection, primary acinar cells were isolated as described previously^23^ and then allowed to recover in oxygenated medium for 2 hours. Total RNA was extracted from fresh isolated acini and subjected to RNA-seq analysis. The extent of *Prdm3* deletion was confirmed in *Prdm3^ΔAcinar^* pancreata **(Supplementary Fig. 5a)**. Analysis of RNA-seq data sets was performed by DESeq2. Setting a *p*-value threshold of 0.05, we identified 1483 genes that were significantly differential expressed (log_2_ |Fold change| > 1) in *Prdm3*-deficient acini compared with control **(Supplementary Table 2)**. Moreover, 1073 out of 1483 differentially expressed genes (DEGs) were upregulated, while only 410 of them were downregulated (**Fig. 5a**). We found that the expression of many adhesion molecules involved in the inflammatory response, such as the TNF receptor superfamily *Tnfrsf21* and *Tnfrsf23*, interleukin receptors *Il18r1* and *Il1r2*, and toll like receptors *Tlr1*, *Tlr2* and *Tlr3* were significantly altered in *Prdm3*-deficient cells. Gene Set Enrichment Analysis (GSEA) further demonstrated that genes involved in inflammatory response and regulation of NF-κB signaling were significantly enriched in *Prdm3*-deficient acinar cells **(Fig. 5b)**. This suggests that loss of *Prdm3* changes cellular homeostasis and favors acinar cells toward a proinflammatory state.

**Figure 5.**
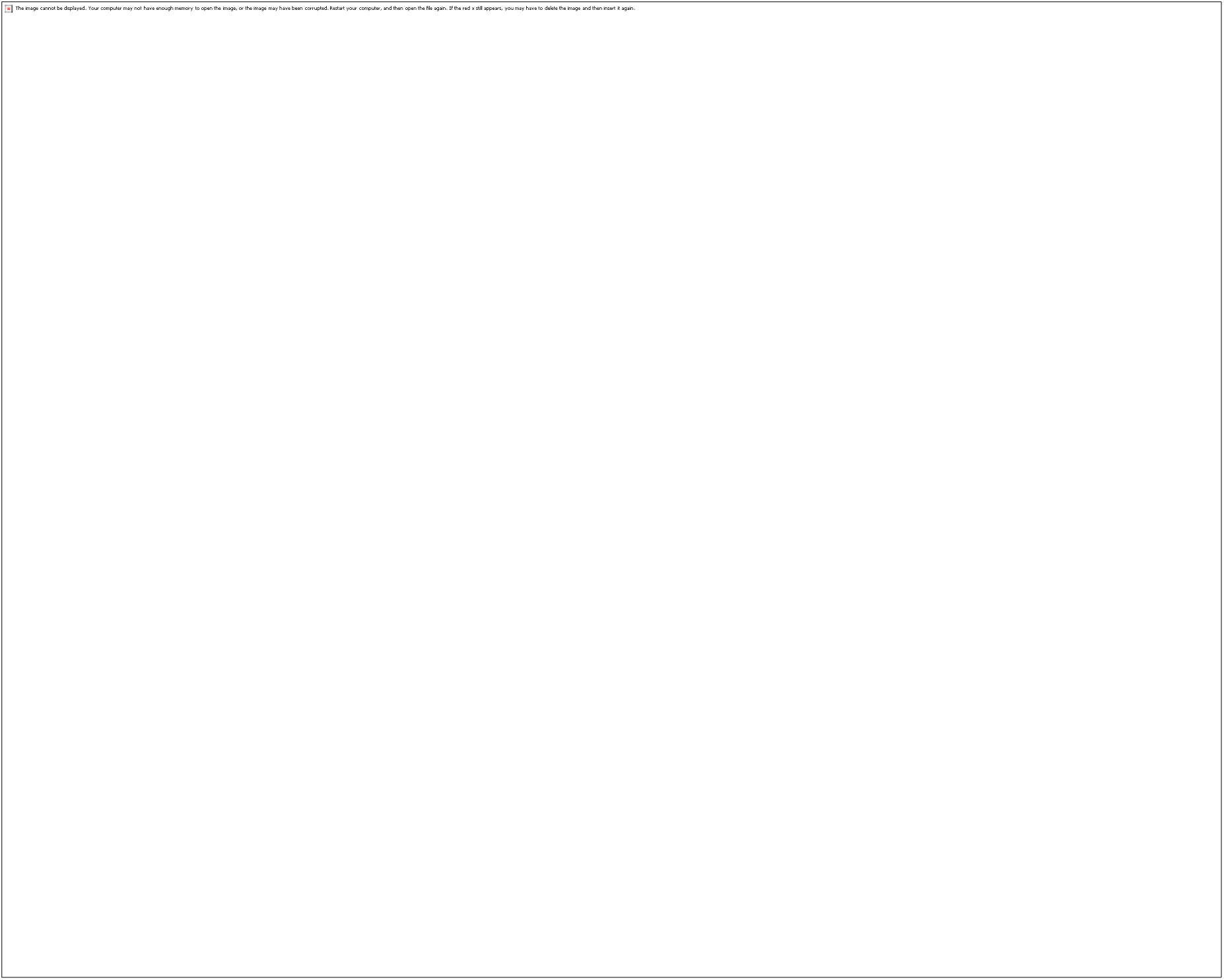
Inactivation of Prdm3 enhances inflammatory response pathways. **(a)** Volcano plot showing a total of 1483 differentially expressed genes in primary acinar cells isolated from *Prdm3^ΔAcinar^* mice, relative to control *Ptf1a^CreER^*. Individual genes are labeled and circled in black. Volcano plot showing differentially expressed genes belonging to inflammatory responses and NF-κB signaling are highlighted in red and blue, respectively. Gene Set Enrichment Analysis (GSEA) of differentially expressed genes (*Ptf1a^CreER^* vs *Prdm3^ΔAcinar^*) identified enrichment of immune responses and regulation of NF-κB signaling. Normalized enrichment score (NES) and *p* values are shown. **(c)** Functional annotation on 1073 upregulated genes in primary acinar cells isolated from *Prdm3^ΔAcinar^* mice, relative to control. Significant KEGG and Hallmark terms, *p*-values and ranks are shown.

To further identify additional biological pathways that were activated upon *Prdm3* deletion, we performed functional annotation on the 1703 upregulated genes with KEGG and Hallmark data sets. Loss of *Prdm3* significantly affected over 30 pathways **(Supplementary Table 3)** including the Hif-1 signaling pathways **(Fig. 5c)**. Intriguingly, recent studies demonstrated that mice with acinar cell-specific deletion of *Hif1*α were less susceptible to cerulein-induced pancreatitis^24^. As our transcriptome analysis revealed a significant elevation of *Hif1a* in *Prdm3*-deficient cells, we compared the protein level of Hif1a in control and *Prdm3^ΔAcinar^* pancreata. We confirmed that the expression level of *Hif1a* was significantly upregulated in *Prdm3*-depleted pancreatic tissue by immunoblotting and immunohistochemical staining **(Supplementary Fig. 5**b-5c). Our findings are supported by a previous study in which shRNA knockdown of *Prdm3* upregulated *Hif1a* in DA-1 and NFS-60 leukemic cells^25^. As loss of *Prdm3* exaggerated inflammation, we speculate that dysregulation of *Hif1a* expression might contribute to the dramatic effects of *Prdm3* depletion on pancreatitis.

## Discussion

In the present study, we investigated the role of *Prdm3* in pancreatitis and pancreatic tumorigenesis using mouse models to indelibly delete *Prdm3* in adult acinar cells. We demonstrated that PRDM3 was substantially upregulated in pancreatic acinar cells of patients with pancreatitis, as well as well-differentiated PDAC, but not poorly differentiated PDAC. Interestingly, our clinical relevance analysis suggests a prognostic relevance of PRDM3 underexpression in PDAC patients with surgical resection. Consistent herewith, we further demonstrated that loss of *Prdm3* not only increased the severity of cerulein-induced pancreatitis, but also accelerated cellular atypia and tumorigenic potential in the pancreas, as *Prdm3*-deficient mice undergo robust formation of precursor lesions in the presence of oncogenic *Kras*. These findings implicate a potentially protective mechanism of *Prdm3* in pancreatic exocrine cells, which is different from the pro-tumor role of PRDM3 in several aggressive forms of cancer including colon, breast and ovarian cancer^26–28^. A number of alternatively spliced variants, including the long and short forms of PRDM3, are expressed in pancreatic cancer cells^14^. Our studies cannot distinguish which forms of PRDM3 are necessary for its anti-tumor effect in acinar cells, therefore, more work will be needed to examine the effects of different PRDM3 isoforms on pancreatitis and pancreatic cancer initiation.

Previous studies demonstrated that Prdm3 promotes proliferation and migration in established pancreatic cancer cell lines^14^. However, when *Prdm3* was knocked out in pancreatic acinar cells, we observed that *Prdm3* depletion potentiated pancreatic cancer initiation and progression to high-grade lesions including ductal carcinoma *in situ*, suggesting that Prdm3 plays a suppressive role in acinar-to-ductal transformation. At a molecular level, we found that over 70% of the differentially expressed genes between normal and *Prdm3*-deficient acinar cells were upregulated in *Prdm3^ΔAcinar^* mice, which is consistent with previous studies demonstrating that Prdm3 acts as a transcriptional repressor through interaction with a variety of co-repressors, such as CtBP, histone methyltransferase SUV39H1 and deacetylase HDAC1/2^10^. We therefore postulate that increased Prdm3 expression in transformed ductal-like cells plays an inhibitory role in pancreatic tumorigenesis through the ability of Prdm3 to suppress a series of signaling cascades important for malignant transformation in exocrine pancreas.

Here we showed that *Prdm3*-deficient acinar cells were much more susceptible to cerulein-induced injury. Loss of *Prdm3* enhanced ADM formation and immune cell activation after cerulein treatment. Our transcriptome analysis suggests that deletion of *Prdm3* in adult pancreatic acini induced significant changes in expression for genes involved in inflammatory response and regulation of NF-κB signaling which is constitutively activated in pancreatitis and pancreatic cancer^29^. Several studies have suggested that inflammation leads to an increase of Ras activity and amplifying oncogenic Ras signaling is necessary for pancreatic cancer progression^30–32^. Activation of Kras alone in mice leads to PDAC formation at a low frequency and takes over one year^33^. In contrast, when pancreatitis was induced in *Kras^G12D^* mice, tumorigenesis occurred within a few months^8, 34^ suggesting that precancerous lesions can arise from acinar cells through a process dramatically hastened by inflammation^35^. In this study, we demonstrated that loss of *Prdm3* accelerated *Kras^G12D^*-induced PanIN initiation and promoted rapid progression of pre-neoplastic lesions to invasive lesions. Given that activation of inflammatory response elevates constitutive Ras activity, we propose that, loss of *Prdm3* upregulates the expression of genes involved in inflammatory response in pancreatic acinar cells, which modulates acinar cell homeostasis to increase sensitivity of acinar cells to inflammatory stimuli and promote widespread formation of precancerous lesions among Kras mutated acinar cells.

Additionally, we found that hypoxia inducible factor *Hif1a* was significantly upregulated in *Prdm3*-deficent acinar cells at both mRNA and protein levels. Our findings are consistent with the previous study in which knockdown of Prdm3 upregulated *Hif1a* in DA-1 and NFS-60 leukemic cells^25^. It has also recently been established that HIF-1 signaling plays important roles in both pancreatitis and pancreatic cancer. *Hif1a* is overexpressed in chronic pancreatitis^36^ and its high expression is associated with poor prognosis in PDAC^37, 38^. Acinar-cell-specific deletion of *Hif1a* prevented intrapancreatic coagulation of fibrinogen and protected mice from cerulein-induced acute pancreatitis^24^, suggesting a functional role of Hif1a in the development of pancreatitis. Given that loss of *Prdm3* exaggerated inflammation, we speculate that dysregulation of *Hif1a* expression might contribute to the dramatic effects of *Prdm3* depletion on pancreatitis.

Taken together, our data demonstrated that loss of *Prdm3* not only increased the severity of cerulein-induced pancreatitis, but also accelerated cellular atypia and tumorigenic potential in the pancreas, as *Prdm3*-deficient mice undergo robust formation of precursor lesions in the presence of oncogenic *Kras*. We uncovered a previously unappreciated role for Prdm3 as a suppressor of both pancreatitis and pancreatic tumorigenesis presumably through regulating inflammatory and Hif-1 signaling pathways in the pancreatic acinar cells.

## Materials and methods

### Human samples

A total of 94 patients diagnosed with PDAC and 22 patients diagnosed with chronic pancreatitis between 2003-2011 were included in this study in accordance with institutional guidelines and approved by the Clinical Research Ethics Committee of the First Affiliated Hospital at Sun Yat-sen University. Written informed consent was received from participants prior to inclusion in this study.

### Mice

*Mice.* All animal experiments were approved by the University of Macau Animal Ethics Committees and carried out in accordance to recommendations stated in the Guide for the Care and Use of Laboratory Animals for the National Institutes of Health (US Department of Health, Education, and Welfare). *Prdm3^flox/flox^*, *Kras^LSL-G12D^*, and *Ptf1a^CreER^* mice have previously been described. Recombination was induced by four subcutaneous injection of tamoxifen every other day at 125 mg/kg body weight on animals at 4 to 5 weeks of age.

### Cerulein-induced Pancreatitis

To induce acute pancreatitis, experimental mice were fasted overnight before administration of cerulein as described previously^4^. Cerulein (American Peptide, Sunnyvale, CA) was dissolved in saline and administrated intraperitoneally at 50 μg/kg body weight hourly for 8 hours. Mice were sacrificed after 3 hours recovery. Alternatively, mice were injected with cerulein (50 μg/kg body weight) at hourly intervals for 8 hours per day for two consecutive days and sacrificed at 48 hours after the last injection. To accelerate tumorigenesis, mice were given hourly injections of cerulein (50 μg/kg body weight) for 6 hours per day on alternating days separated by 24 hours and sacrificed after 21 days.

### Histology and immunohistochemical analyses

Paraffin-embedded sections were prepared and subjected to hematoxylin, eosin Y, Alcain Blue or immunohistochemical staining. H&E staining and IHC followed our established procedures^5^, including antigen retrieval with citrate buffer (pH 6.0) prior to staining paraffin sections. For Alcian Blue staining, paraffin sections were incubated in 3% acetic acid for 3 minutes, followed by staining in 1% Alcian Blue staining solution for 30 minutes, and subsequently in Nuclear Fast Red for 5 minutes. All slides were scanned with a 20× objective using a 2D glass slide digital scanner (Leica Biosystems, Vista, CA) and examined at high magnification using the Aperio ImageScope software (Leica). The Aperio positive pixel Algorithm was used to quantify area with positive staining and Aperio nuclear V9 algorithm was used to quantify the number of nuclei. The percentages of Cpa1-, CK19-, Muc5AC-, and Alcian Blue-positive area were calculated by positive pixels divided by the total pixels in selected tissue areas. The percentages of Ly6B.2-, and Prdm3-positive cells were calculated by positive number of nuclei divided by the total number of nuclei in selected tissue areas. Six sections, which displayed maximal pancreatic cross-sectional area, from each animal were used for quantification. The number of PanINs were counted based on the characteristics of every gland with cuboidal to columnar duct-like structures and enlarged lumen in six sections per mouse. For the number of high-grade lesions, including PanINs and ductal carcinoma *in situ*, five 1600 × 920 μm squares in one section per mouse were analyzed and the squares were randomly distributed in the pancreas. Primary and secondary antibodies used for staining is provided in Supplementary Table 4.

### Evaluation of immunostaining of PRDM3 in patient specimens

PRDM3 expression was evaluated according to the staining intensity and proportion of positively stained tumor cells. Staining intensity was graded as 0 (negative), 1 (weakly positive), 2 (moderately positive), and 3 (strong positively). The proportion of positively stained tumor cells was scored as 0 (no positive cells), 1 (< 10% positive cells), 2 (10 - 25% positive cells), 3 (25 - 50% positive cells), and 4 (> 50% positive cells). The immunostaining of PRDM3 was determined by staining index (SI) through multiplying the staining intensity by the proportion of positively stained tumor cells as previously described^39^. The expression levels of PRDM3 was regarded as high if the SI score is > 6, or low if the SI score if ≤ 6. The immunohistochemical specimens were evaluated by two independently pathologists who were blinded to clinical diagnosis.

### RNA isolation and quantitative real-time PCR

Mice were sacrificed with CO_2_ asphyxiation followed by cervical dislocation. Pancreata were immediately harvested and cut into small pieces in RNALater (Qiagen, Redwood, CA). To detect mRNAs, twenty mg of tissue was homogenized in 1 ml of Trizol using a T 10 basic ULTRA-TURRAX® homogenizer. RNA was extracted from Trizol according to the manufacturer’s instructions (Thermo Fisher Scientific, Waltham, MA) and subsequently subjected to reverse transcription using PrimeScript™ RT reagent Kit (TaKara Bio, Mountain View, CA). Quantitative real-time PCR was performed with Premix Ex Taq (TaKara Bio) on CFX96 qPCR system (BioRad Laboratories). Results were normalized to *Gapdh* for mRNA detection. The quantitative real-time PCR primer sequences are list in Supplementary Table 5.

### Western blotting

Pancreata were harvested and snap frozen in liquid nitrogen. The frozen tissue was homogenized in lysis buffer containing 20 mM Tris-HCl (pH 7.5). 150 mM NaCl, 1 mM Na_2_EDTA, 2mM Na_2_VO_4_, 1% Triton X-100, 5mM 4-nitrophenyl phosphate, 0.5% sodium deoxylcholate, 1mM phenylmethanesulfonylfluoride, protease inhibitor cocktail (Sigma). Twenty μg of protein was separated on a 10% sodium dodecyl sulfate-polyacrylamide gel, transferred onto a polyvinylidene difluoride membrane, and probed with antibodies. The membranes were visualized using ECL™ Western Blotting Detection System (GE Healthcare) and ChemiDoc™ Imaging Systems (BioRad Laboratories).

### Serum amylase assay

Blood was collected by cardiac puncture and placed at room temperature for 30 minutes. Serum was separated from red blood cells by centrifugation at 2500 g for 15 min. The top layer which contained serum was transferred to a new tube for further analysis. Amylase activity was measured using the Amylase Colorimetric Assay Kit following manufacturer’s instructions (Sigma).

### Isolation of primary acinar cells

Primary acinar cells were isolated as described in detail previously^23^ with a small modification. In brief, pancreas was harvested and transferred into ice-cold Hank’s balanced salt solution (HBSS). Lymph nodes, fat and mesenteric tissues were carefully removed. Pancreas was minced into 2- to 3-mm pieces and digested with 0.2 mg/ml collagenase P (Roche) at 37 °C for 10-12 minutes. Cell clusters were washed 3 times with ice-cold HBSS containing 5% FBS and filtered through 100-μm cell strainer (BD Biosciences). The cell suspension containing acini was carefully layered on top of HBSS containing 30% FBS. Primary acinar cells were pelleted (80 g, 2 min, at 4 °C) and re-suspended in Waymouth media (Sigma). To obtain acinar-cell-conditioned media, primary acinar cells were incubated in Waymouth media (10% FBS and 10 mM HEPES) containing 10 nM cerulein for 6 hours. Supernatants were collected for macrophage activation experiment.

### RAW246.7 cell culture and activation

Raw264.7 macrophage cells were obtained from American Type Culture Collection (ATCC) and maintained in DMEM containing 10% FBS and 100 U/ml penicillin/streptomycin in a 37 °C humified incubator supplemented with 5% CO_2_. To stimulate macrophage activation, 5 × 10^5^ Raw264.7 cells per well of 24-well plate were culture in DMEM (10% FBS and 100 U/ml penicillin/streptomycin) overnight and then incubated in acinar-cell-conditioned media for 16 hours. RNA was extracted for quantitative real-time PCR. Data were normalized to *Gapdh* and the number of primary acinar cells used to prepare conditioned media.

### RNA-seq analyses

Total mRNA was extracted from primary acinar cells freshly isolated from 6-week-old *Ptf1a^CreER^;Prdm3^flox/flox^*mice and their corollary controls (*Ptf1a^CreER^*), one week after the last tamoxifen injection (125 mg/kg, 4 times, every other day). RNA concentration and integrity were measured using the Agilent 2100 Bioanalyzer (Agilent Technologies). cDNA libraries were prepared using NEBNext® Ultra™ RNA Library Prep Kit for Illumina (New England Biolabs) according to the manufacturer’s instructions. Libraries were sequenced at Novogene (Tianjin, China) with 100× coverage and 150 bp paired end reads on Illumina HiSeq 2500 instrument. The quality of the sequencing data was analyzed by using FastQC (version 0.11.5), and raw reads with low quality were removed using Trim Galore (version 0.4.4) prior to analysis of the data. All the trimmed reads were mapped to reference mouse genome (mm10, GRCm38) by using STAR (version 020201), and the mapped counts were extracted using feature count from Subread package (version 1.5.3). Subsequently, read count data containing 49,492 quantified transcripts with raw reads was preprocessed by filtering out genes with zero read count across different samples, and 20,069 genes remained after filtering. The read count data were normalized by DESeq2, which can be used for downstream differential expression analysis. Genes differentially expressed (DEG) in *Prdm3^ΔAcinar^* vs control were filtered by log2 fold change > 1 or < -1. P value < 0.05 was considered to be significantly different. Gene set enrichment analysis (GSEA) was performed using Bioconductor R package clusterProfiler^40, 41^. The gene sets implemented were derived from Cancer hallmark, Kyoto Encyclopedia of Genes and Genomes (KEGG) and Gene Ontology (GO), which was collected in the Molecular Signatures Database (MSigDB; version 6.2). Functional enrichment was performed on genes up-regulated *Prdm3*-deficient cells.

### Statistical Analysis

All data are presented as mean ± SD from at least three mice in each experimental group. Statistical analysis of different animal groups was acquired using the t test with GraphPad software. Statistical analyses of patient samples were performed with SPSS software using two-sided t test. Kaplan-Meier survival plots were generated using the log-rank test. *p* value < 0.05 was considered statically significant.

## Acknowledgements

We thank the University of Macau, Faculty of Health Sciences, Animal Research Facility for animal housing. We thank Christopher Wright (Vanderbilt University) for *Ptf1a^CreER^* mice and David Tuveson (Cold Spring Harbor Laboratory) for *LSL-Kras^G12D^*mice. This work was supported by the National Natural Science Foundation of China (NSFC 31701276) and the Science and Technology Development Fund of Macau SAR (FDCT 170/2017/A3 and FDCT 0033/2019/A1) to R.X, Canadian Institutes of Health Research Operating Grant (mop-142216) and New Investigator Award to J.L.K., National Natural Science Foundation of China (NSFC 81672417) to W.L., and National Institutes of Health (R01-DK078803) to M.S..

## Supplementary materials

**Supplementary Fig. 1.**
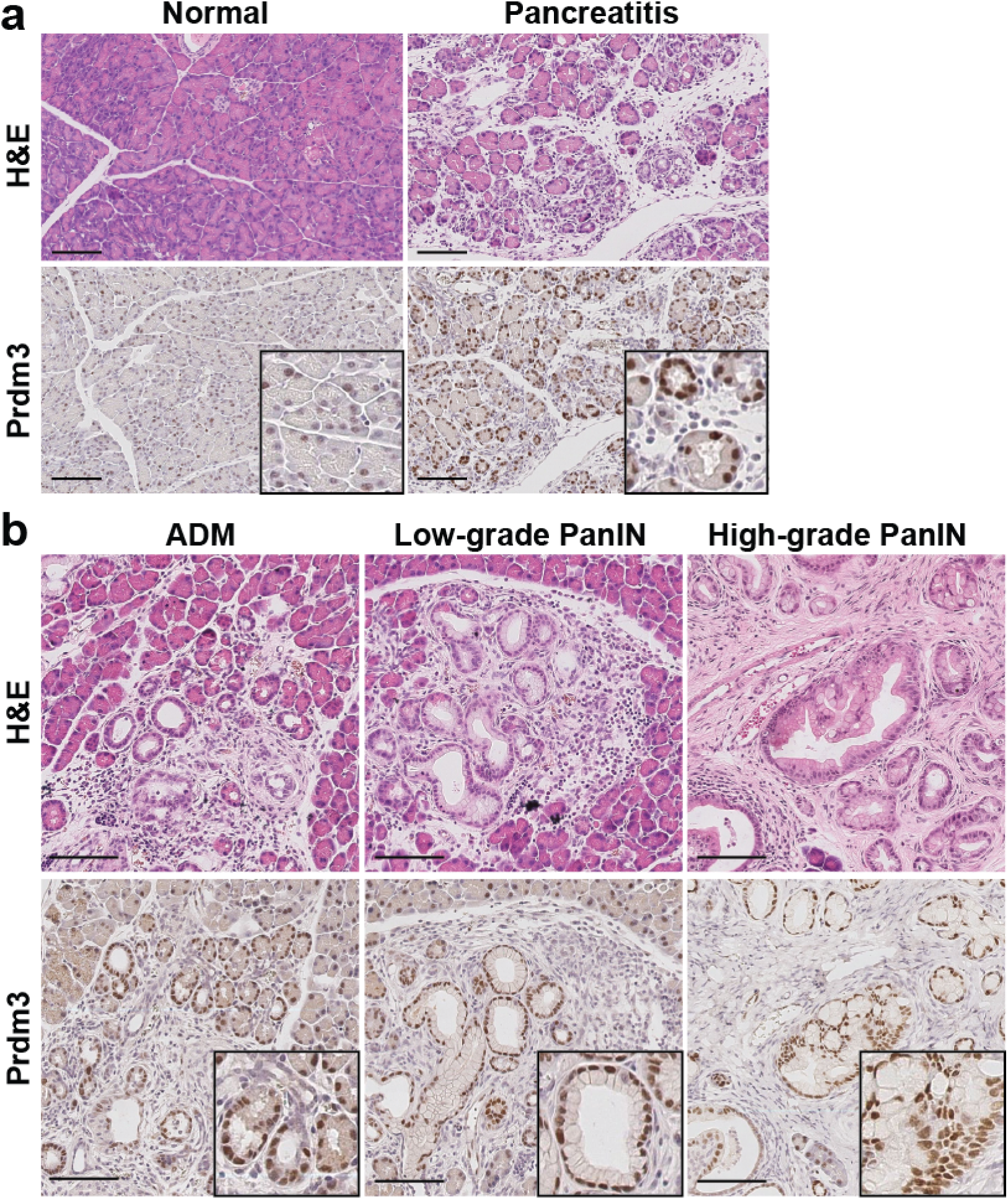
Elevated PRDM3 expression is associated with inflamed pancreatic epithelia as well as premalignant pancreatic malignant lesions. **(a)** C57BL/6 mice were injected with cerulein (50 μg/kg, 8 hourly per day, 2 consecutive days) to induce acute pancreatitis. Pancreata were harvested 48 hours after the last cerulein injection and subjected for hematoxylin-eosin staining (H&E) and Prdm3 immunostaining. **(b)** Immunohistochemistry for Prdm3 and H&E of ADM, low-grade PanIN and high-grade PanIN of *Ptf1a^CreER^;Kras^G12D^* pancreata. Scale: 100 μm

**Supplementary Fig. 2.**
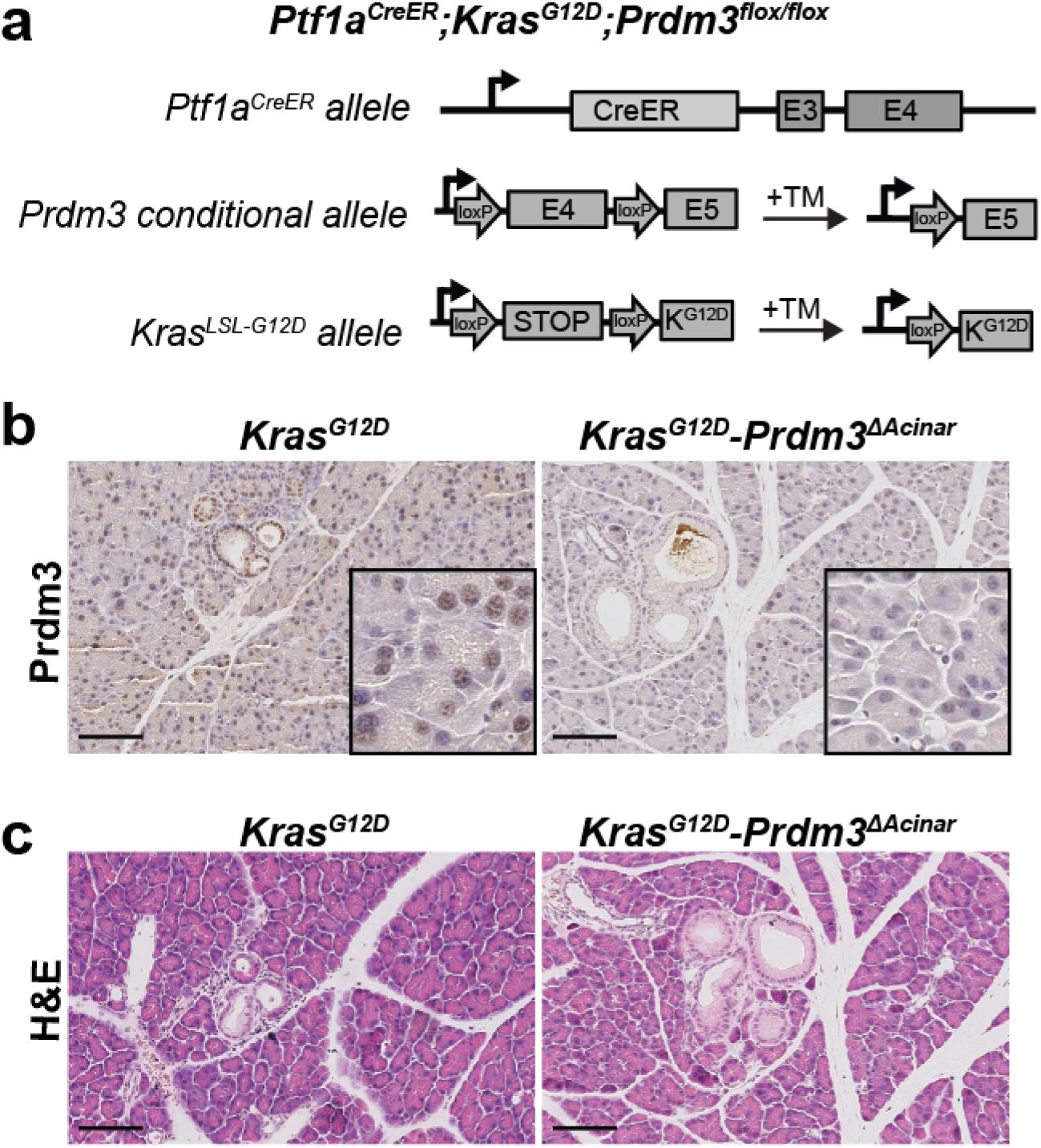
Generation of Prdm3 conditional knockout mouse. **(a)** Strategy to generate Prdm3 conditional knockout mice: *Ptf1a^CreER^;Kras^G12D^;Prdm3^flox/flox^*mice. Mice were injected with tamoxifen TM at 4-5 weeks of age and pancreata were analyzed at 4 weeks after the last tamoxifen injection. Immunohistochemistry for Prdm3 **(b)** and Hematoxylin-eosin staining (H&E) **(c)** in *Kras^G12D^-Prdm3^ΔAcinar^* and *Ptf1a^CreER^* control mice. Scale: 100 μm.

**Supplementary Fig. 3.**
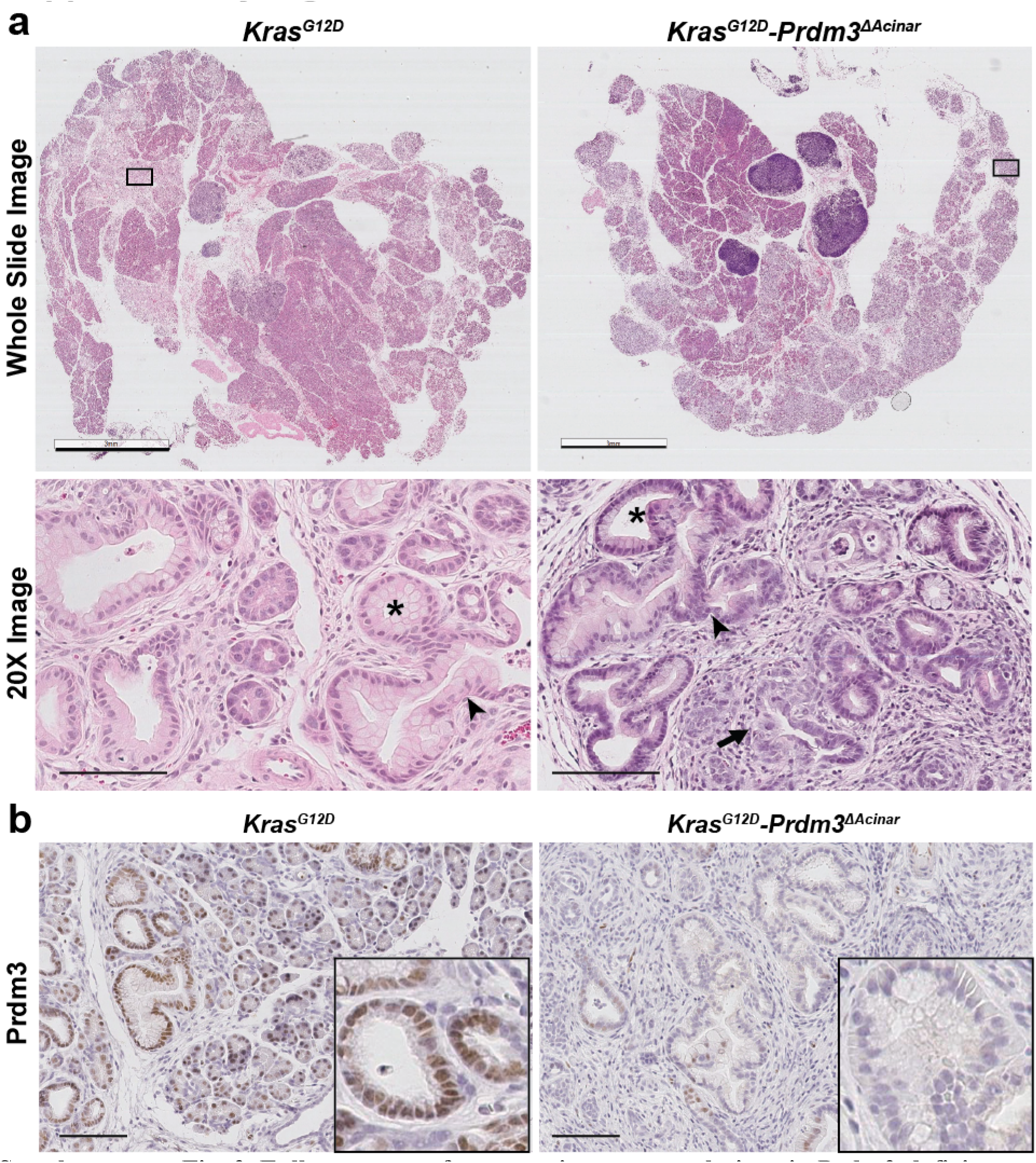
Full spectrum of pancreatic precursor lesions in Prdm3-deficient mice. **(a)** Hematoxylin-eosin staining of representative whole-section and high-magnification images of *Kras^G12D^-Prdm3^ΔAcinar^* and *Kras^G12D^* pancreata 21 days post-cerulein injection. Low-grade PanINs are indicated by stars, high-grade PanINs are indicated by arrowheads, and ductal carcinoma *in situ* is indicated by arrow. **(b)** Immunohistochemistry for Prdm3 in *Kras^G12D^-Prdm3^ΔAcinar^* and *Kras^G12D^* mice. Scale: 3 mm (a top) and 100 μm (a bottom and b).

**Supplementary Fig. 4.**
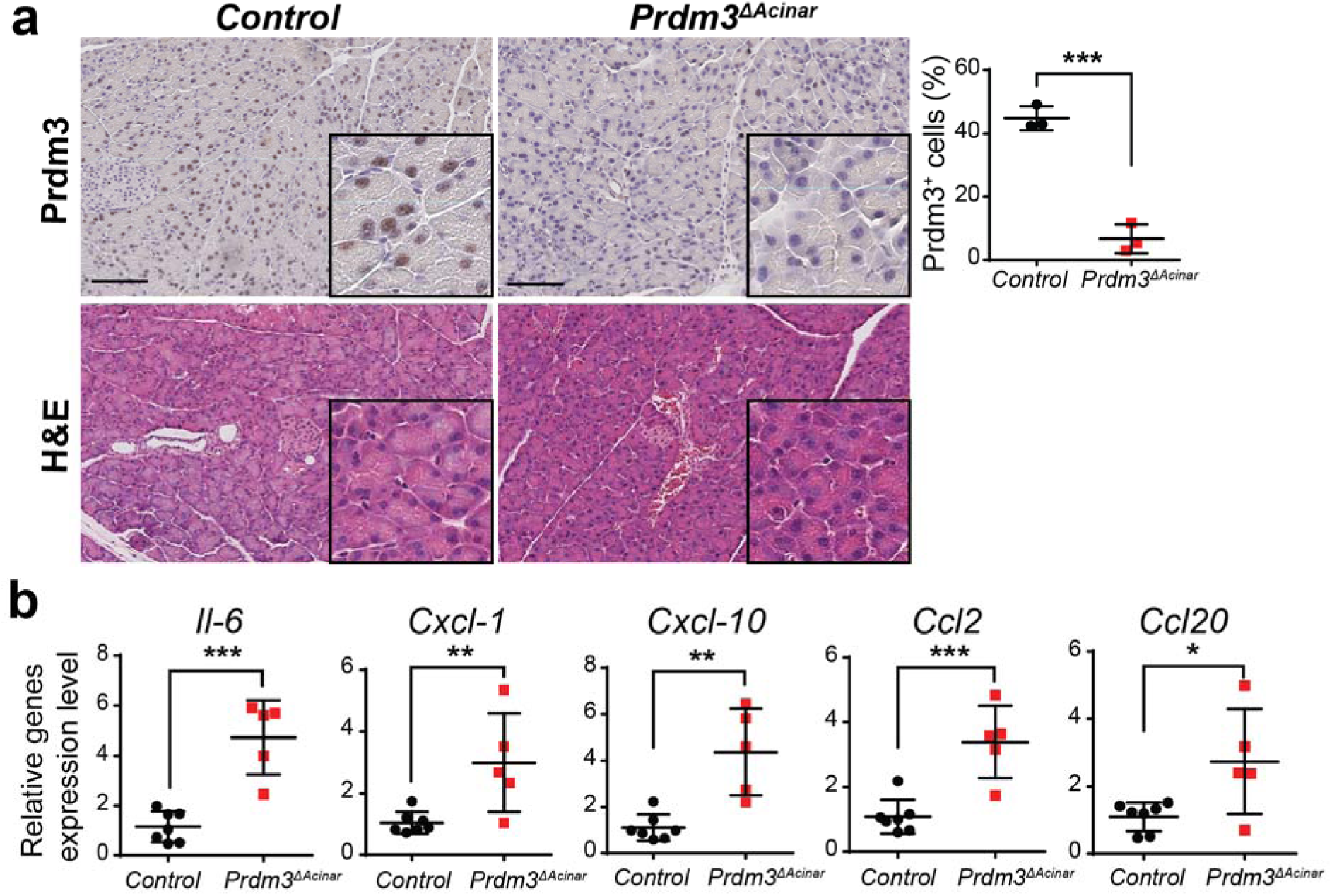
Prdm3-deleted mice are more susceptible to cerulein-induced inflammatory responses. **(a)** *Ptf1a^CreER^* control and *Ptf1a^CreER^;Prdm3^flox/flox^*mice were injected with tamoxifen at 4-5 weeks of age and pancreata were analyzed at 7 days after the last tamoxifen injection. Hematoxylin-eosin staining (H&E) and immunohistochemistry for Prdm3. **(b)** Quantitative real-time PCR for indicated cytokines in control (n=7) vs *Prdm3^ΔAcinar^* (n=5) pancreata from mice 3 hours after injection of cerulein 8 times at hourly intervals. Data show mean ± SD. \**p* < 0.05, \*\**p* < 0.01, *** *p* < 0.001. Scale: 100 μm.

**Supplementary Fig. 5.**
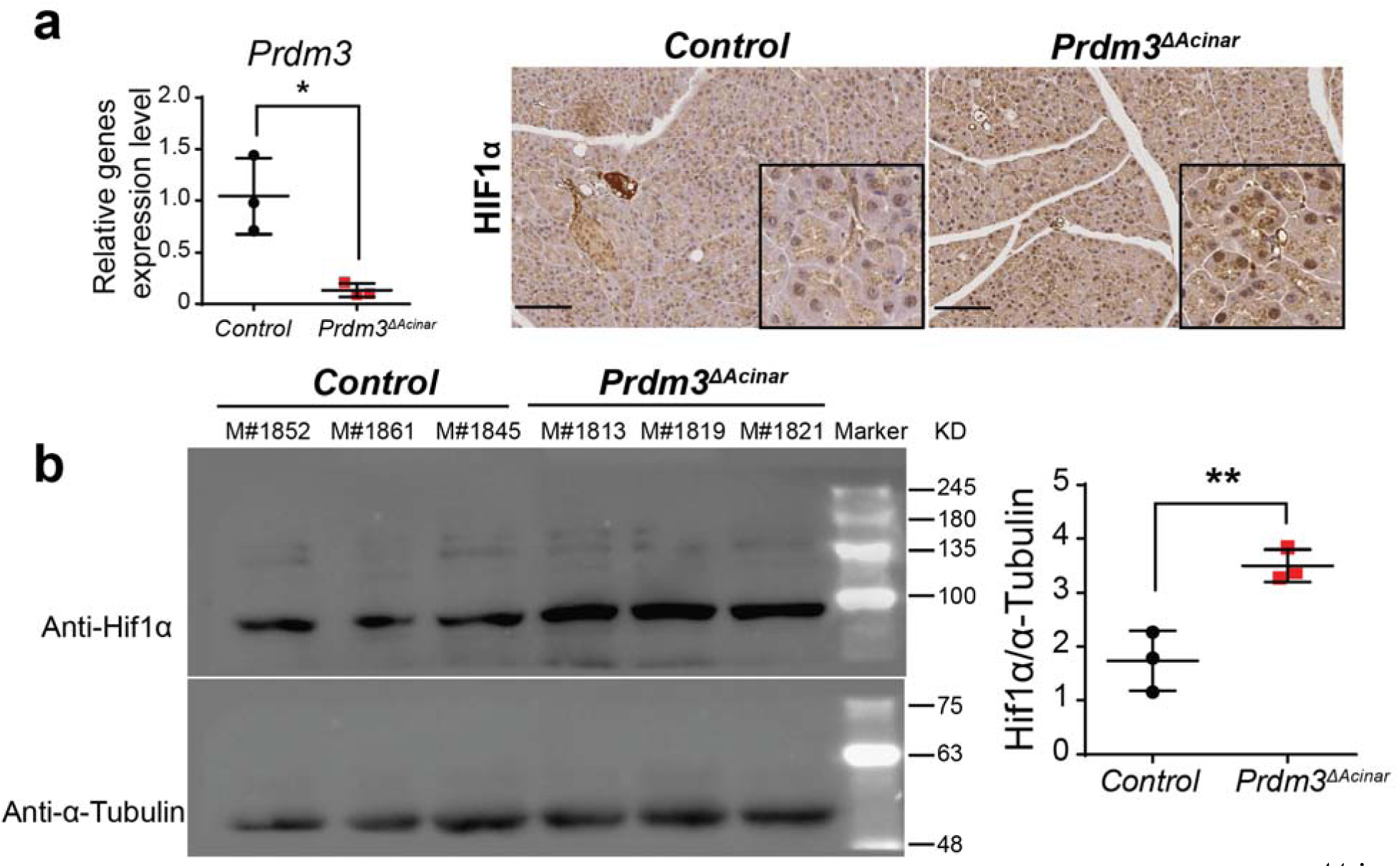
Immunostaining reveals an upregulation of Hif1α in *Prdm3^ΔAcinar^* pancreata. **(a)** Quantitative real-time PCR for *Prdm3* in primary acinar cells isolated from *Prdm3^ΔAcinar^* (n=3) and control pancreata (n=3). **(b)** Hif1α immunoblot analysis of pancreas lysates from *Ptf1a^CreER^* control and *Prdm3^ΔAcinar^* mice. Equal loading was confirmed by immunoblot with an anti-α-tubulin antibody. Densitometric analysis of Hif1α normalized to α-tubulin is shown on the right. **(c)** Immunohistochemistry for Hif1α of *Prdm3^ΔAcinar^* and *Ptf1a^CreER^* control pancreata. Data show mean ± SD. \**p* < 0.05, \**p* <0.01. Scale: 100 μm

**Supplementary Table 1.** **PRDM3 expression in normal and pancreatitis tissue**

| PRDM3 expression in normal and pancreatitis tissue |  |
| --- | --- |
| Histology | Staining intensity score (0 / 1 / 2 / 3) |
| Normal (n=8) | 1 / 6 / 1 / 0 |
| Pancreatitis (n=22) | 0 / 3 / 4 / 15 |

**Supplementary Table 2.** Excel spreadsheet indicates differential expressed genes (| Log_2_FC | > 1, *p* < 0.05) in Prdm3-depleted pancreatic acinar cells.

**Supplementary Table 3.** Hallmark and KEGG pathway analyses of upregulated genes in Prdm3-depleted pancreatic acinar cells. Excel spreadsheet shows the output pathways, *p*-values, gene ratio, relative gene counts and gene IDs.

**Supplementary Table 4.** **List of primary and secondary antibodies.**

| Primary antibody |  |  |  |
| --- | --- | --- | --- |
| Antigen | Species | Source | Dilution |
| Prdm3 | Rabbit | Cell signaling | 1:500 |
| Ly6B.2 | Rat | Serotec | 1:1000 |
| Cytokeratin 19 | Rabbit | Abcam | 1:2000 |
| Muc5AC | Mouse | Thermo Scientific | 1:500 |
| Cpa1 | Goat | R&D | 1:1000 |
| Hif1α | Rabbit | NOVUS | 1:500 |
| α-Tubulin | Mouse | Santa Cruz | 1:1000 |
| Secondary antibody | Conjugation | Source | Dilution |
| Rabbit/Rat/Mouse/Goat | Biotinylated | Vector Laboratories | 1:500 |
| Rabbit/Mouse | HRP | Jackson Immunoresearch | 1:2000 |

**Supplementary Table 5.**
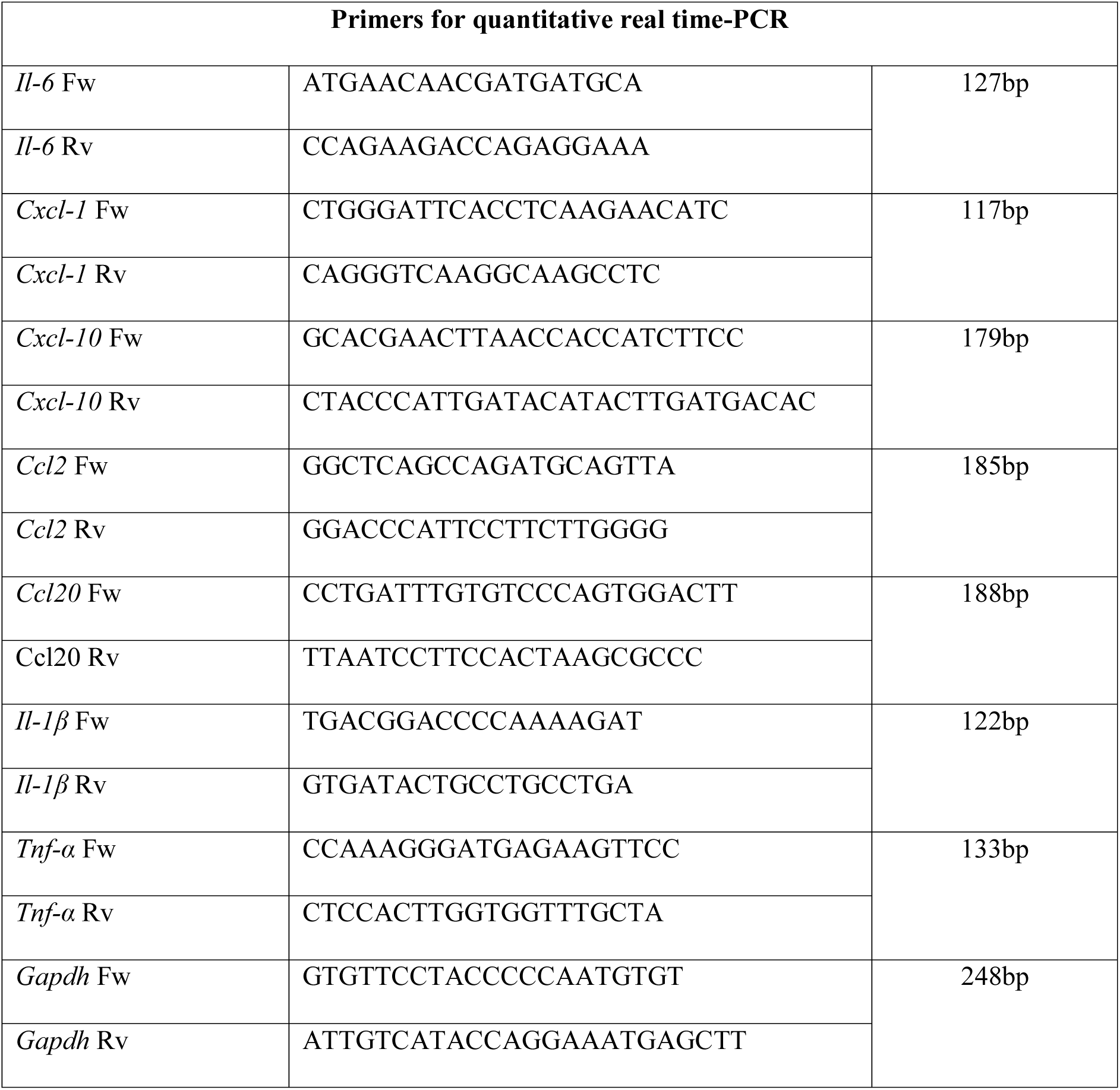
**List of primers for quantitative real time-PCR.**

